# Plakoglobin regulates adipocyte differentiation independently of the Wnt/β-catenin signalling pathway

**DOI:** 10.1101/2022.02.03.478896

**Authors:** F Abou Azar, Y Mugabo, S Yuen, S Allali, S Del Veliz, G Lavoie, PP Roux, GE Lim

## Abstract

The scaffold protein 14-3-3ζ is an established regulator of adipogenesis and postnatal adiposity. We and others have demonstrated that the 14-3-3ζ interactome to be diverse and dynamic, and it can be examined to identify novel regulators of physiological processes, including adipogenesis. In the present study, we sought to determine if factors that influence adipogenesis could be identified in the 14-3-3ζ interactome found in white adipose tissue of lean or obese TAP-tagged-14-3-3ζ overexpressing mice. Using mass spectrometry, changes in the abundance of novel, as well as established, adipogenic factors within the 14-3-3ζ interactome could detected. One novel candidate is plakoglobin, the homolog of the known adipogenic inhibitor β-catenin, and herein, we report that plakoglobin is involved in adipocyte differentiation. Plakoglobin is expressed in murine 3T3-L1 cells and is primarily localized to the nucleus, where its abundance decreases during adipogenesis. Ectopic overexpression and siRNA-mediated depletion of plakoglobin had dual effects on inhibiting adipogenesis and reducing PPARγ2 expression. Plakoglobin depletion in human adipose-derived stem cells also impaired adipogenesis and reduced lipid accumulation post-differentiation. Transcriptional assays indicated that plakoglobin does not participate in Wnt/β-catenin signaling, as its depletion did not affect Wnt3a-mediated SUPERTOPFlash activity. Taken together, our results establish plakoglobin as a novel regulator of adipogenesis *in vitro* and highlights the ability of using the 14-3-3ζ interactome to discover undiscovered pro-obesogenic factors.

## Introduction

White adipose tissue primarily functions as a storage depot for nutrients and triglycerides and influences metabolism through the secretion of hormones and cytokines (1,2). The expansion of fat mass occurs by hyperplasia of adipocytes during the early stages of development and through hypertrophy during adulthood (3,4). Hypertrophy refers to the increase in size of adipocytes, while hyperplasia is characterized by increases in adipocyte number (5,6). Murine adipogenesis occurs via a two-step process, where the first phase involves mesenchymal stem cells committing to the adipocyte lineage and differentiating into pre-adipocytes. The second step is defined by pre-adipocytes differentiating into mature adipocytes, and this requires the expression of the master adipogenic regulator, PPARγ2 (2). While the sequential and parallel mechanisms responsible for adipogenesis have been extensively studied, our knowledge of the factors that can influence the process remain incomplete.

Previous studies have highlighted the inhibitory effects of the Wnt/β-catenin signalling pathway on adipogenesis, and activation of the Wnt signaling cascade promotes mesenchymal stem cells to differentiate to osteoblasts and blocks the commitment towards the adipocyte lineage (7–11). Differences in the effect of inhibiting Wnt signaling on adipogenesis have been observed *in vitro* and *in vivo*. In pre-adipocyte cell lines, inhibition of the Wnt/β-catenin pathway permits spontaneous adipocyte differentiation (10). However, it has been recently shown that deletion of β-catenin in primary mouse adipocytes is associated with resistance to high-fat diet-induced obesity due to decreased expansion of subcutaneous adipocytes, decreased proliferation of Pdgfrα+ adipocyte progenitor cells, and loss of adipocyte maturity (12). Taken together, these findings demonstrate the complexities involved in studying Wnt/β-catenin-associated effects that influence adipogenesis.

Plakoglobin, also known as γ-catenin, is a close homolog of β-catenin. They share a high degree of sequence similarity, and both plakoglobin and β-catenin contain 12 Armadillo repeats that allow for interactions with a wide variety of targets and results in overlapping functions (13). Both plakoglobin and β-catenin are required for cell-cell adhesion, contributing to the formation of adherens junctions that confer structural integrity to epithelial and non-epithelial cells (14–16); however, they can also have divergent functions, as only plakoglobin is present in desmosomes where it participates in cell-cell adhesion and provide resistance to mechanical stress (17). Plakoglobin may also compete with β-catenin for binding to TCF7L2, the transcriptional coactivator of the Wnt signalling pathway and inhibit β-catenin-dependent transcriptional activity in a context dependent manner (18). For example, cardiomyocyte-specific overexpression of plakoglobin was found to inhibit the transcriptional activity of the Wnt/β-catenin signaling pathway and promoted adipogenesis (19). However, whether plakoglobin can directly influence adipocyte differentiation has yet to be studied.

Scaffold proteins of the 14-3-3 family are highly conserved and ubiquitously expressed, holding multiple functions related to proliferation, apoptosis, and metabolism (20–22). 14-3-3 proteins typically exert their effects by binding to phosphorylated serine or threonine motifs on binding partners, allowing them to influence their subcellular localization and activities (23). We have previously shown that the 14-3-3ζ isoform is integral to adipogenesis, as systemic deletion led to a reduction of visceral adipose tissue in mice and depletion in 3T3-L1 pre-adipocytes blocked differentiation (24). While we have found that 14-3-3ζ over-expression potentiated age- and diet-induced weight gain (24), it was unclear if this was due to the recruitment of pro-obesogenic factors to the 14-3-3ζ interactome and changes in their cellular functions. Using affinity proteomics, we previously demonstrated the utility of mass spectrometry-based approaches to elucidate the 14-3-3ζ interactome within mouse embryonic fibroblasts overexpressing 14-3-3ζ as a means of identifying novel regulators of adipogenesis (25). We identified an enrichment of RNA splicing factors within the 14-3-3ζ interactome during adipocyte differentiation and reported an essential role of Heterogeneous Nuclear Ribonucleoprotein F (HNRPF) in the generation of the critical adipogenic splice variant, PPARγ2 (25). Whether such an approach could be used to discern changes between the 14-3-3ζ interactome in the context of diet-induced obesity has not been attempted.

In the present study, enriched abundance of plakoglobin within the 14-3-3ζ interactome was detected by proteomics in gonadal adipose tissue of obese, 14-3-3ζ-overexpressing mice, which suggested its involvement in adipogenesis. Depletion of plakoglobin and β-catenin in 3T3-L1 pre-adipocytes and human adipocyte stem cells reduced and potentiated adipogenesis, respectively. Knockdown of plakoglobin did not affect β-catenin transcriptional activity, implying its actions are independent of its closely-related homolog. Overexpression of either plakoglobin or β-catenin led to a marked decrease in PPARγ2 abundance, demonstrating impaired adipogenesis. The ability to determine pro-adipogenic factors in the 14-3-3ζ interactome within adipose tissue of obese mice demonstrates a novel approach to identify potential pro-obesogenic factors that influence fat expansion.

## Materials and Methods

### Proteomic analysis of the 14-3-3ζ interactome

Gonadal white adipose tissues (n=3 per group) were isolated from wildtype mice or transgenic mice over-expressing a TAP-tagged 14-3-3ζ that were fed a low-fat (LFD) or 60% high-fat diet (HFD; Research Diets, New Brunswick, NJ) for 12 weeks (24). Tissues were homogenized in RIPA buffer supplemented with the Halt Protease and Phosphatase inhibitor cocktail (ThermoFisher Scientific, Waltham, MA), and supernatants were collected following centrifugation. Two milligrams of total protein were incubated overnight at 4°C with Pierce Protein G magnetic beads (ThermoFisher), followed by successive washes. Protein complexes were dissociated from beads with 1X Laemmli sample buffer at 95°C for 15 min under non-reducing conditions. Following SDS-PAGE separation and Coomassie staining, bands were excised and digested in gel with trypsin. Samples were reconstituted in formic acid 0.2% and loaded and separated on a homemade reversed-phase column (150 μm i.d. x 150 mm) with a 56-min gradient from 0–40% acetonitrile (0.2% FA) and a 600 nl/min flow rate on an Easy-nLC II (Thermo Fisher Scientific), connected to a Q-Exactive Plus mass spectrometer (Thermo Fisher Scientific). Each full MS spectrum acquired with a 60,000 resolution was followed by 20 MS/MS spectra, where the 12 most abundant multiply charged ions were selected for MS/MS sequencing. Database searches were performed against the SwissProt mouse database (containing 20 347 protein entries) using PEAK Studio (version 8.5) as search engine, with trypsin specificity and three missed cleavage sites allowed. Methionine oxidation and asparagine/glutamine deamidation were set as variable modifications. The fragment mass tolerance was 0.01 Da and the mass window for the precursor was ± 10 ppm. Note that only proteins with at least two peptides identified and protein probability ≥ 0.95 were considered, which corresponds to an estimated protein level FDR of ~0.5%. Protein abundance was compared between LFD and HFD gonadal adipose tissue samples, and proteins with significantly different levels (log2-fold change >0.5, FDR<0.05) were subjected to gene ontology analysis (ShinyGO v0.741) (26) or STRING (27).

### Cell culture

NIH-3T3 and 3T3-L1 pre-adipocyte cells (ZenBio, Durham, North Carolina) were maintained in 25mM glucose DMEM (Life Technologies Corporation, Grand Island, NY), supplemented with 10% newborn calf serum (NBCS) and 1% penicillin/streptomycin (P/S). Differentiation of confluent 3T3-L1 cells was induced by a cocktail of 500 μM IBMX (Sigma-Aldrich, Oakville, Ontario), 500 nM dexamethasone (Sigma-Aldrich) and 172 nM insulin (Sigma-Aldrich) for 48 hours, as previously described (24,25,28), and cells were maintained with 25mM glucose DMEM with 10% FBS, 1% P/S and 172 nM insulin for up to 7 days. Human adipocyte stem cells (hADSCs) (ZenBio) were maintained in DMEM/F-12 (Wisent, Saint-Bruno, Quebec) supplemented with 10% FBS and 1% P/S. Differentiation was induced by a cocktail of 33μM Biotin/17μM panthothenate, 1μM dexamethasone (Sigma-Aldrich), 1μM insulin, 5μM rosiglitazone and 500μM IBMX (Sigma-Aldrich) for one week. hADSCs are maintained for an additional week in induction media without rosiglitazone and IBMX. To assess lipid incorporation, differentiated cells were stained with Oil Red-O (ORO) and absorbance of ORO was quantified, as previously described (24,28).

### Transient transfection of siRNA and plasmids

Knockdown of β-catenin, plakoglobin, or 14-3-3ζ in 3T3-L1 cells was achieved by transfecting preadipocytes one day prior to the induction of differentiation with 10 nM Silencer Select siRNAs (ThermoFisher) targeting *Ctnnb1*(s63417, s63118), *Jup* (s68573, s68571), and *Ywhaz* (s76189) mRNA, respectively. In hADSCs, cells were transfected with 10 nM Silencer Select siRNAs (ThermoFisher) targeting *CTNNB1*(s438, s437), *JUP* (s7666, s534842) and *YWHAZ* (s14972), *YWHAH* (s14968), *YWHAQ* (s21598), *YWHAS* (s5961), *YWHAE* (s200463), *YWHAQ* (s14965), *YWHAB* (s14961), *YWHAZ* (s14972), 2 days prior to differentiation.

To over-express plakoglobin and β-catenin, the plasmids mEmerald-JUP-C-14 and mEmerald-Beta-Catenin-20 were transfected in 3T3-L1 pre-adipocytes by Nucleofection (Lonza). mEmerald-JUP-C-14 (Addgene plasmid # 54132; http://n2t.net/addgene:54132; RRID:Addgene_54132) and mEmerald-Beta-Catenin-20 (Addgene plasmid # 54017; http://n2t.net/addgene:54017; RRID:Addgene_54017) were gifts from Michael Davidson.

To measure Wnt/β-catenin transcriptional activity, NIH 3T3 cells were transfected with M50 Super 8xTOPFlash reporter (Addgene plasmid #12456; http://n2t.net/addgene:12456; RRID:Addgene_12456) or M51 Super 8xFOPFlash reporter (Addgene plasmid #12457; http://n2t.net/addgene:12457; RRID:Addgene_12457)) plasmids, both of which were gifts from Randall Moon, by Lipofectamine 3000 (Thermofisher Scientific). Renilla luciferase was used as internal control. Following transfection with SuperTOPFlash or Super 8xFOPFlash, knockdowns of β-catenin, plakoglobin, and 14-3-3ζ were achieved by transfecting cells with siRNA. Three days after the initial transfection with SuperTOPFlash or SuperFOPFlash, cells were exposed to 40mM of LiCl (Sigma-Aldrich, #L4408-100G) or 0.5ng/mL of Wnt3a recombinant protein (BioLegend, San Diego, CA, #772301). Luciferase activity was measured by bioluminescence (Promega Dual-Glo Luciferase Assay System, Madison WI) on a Victor3V Multilabel Counter (PerkinElmer, Waltham, MA).

### RNA isolation and quantitative real-time PCR

Total RNA was extracted from cells using RNeasy Mini kit (Qiagen). Reverse transcription of RNA to cDNA was performed with the High-Capacity cDNA Reverse Transcription kit (ThermoFisher Scientific). Real-time PCR was carried out on the QuantStudio 6-flex Real-time PCR System (ThermoFisher Scientific) using SYBR Green FastMix (QuantaBio, Beverly, MA). Data were normalized to *Hprt* or *BACTIN* and analyzed using the ΔΔCt method. All primer sequences are in Table 1.

**Table 1-.**
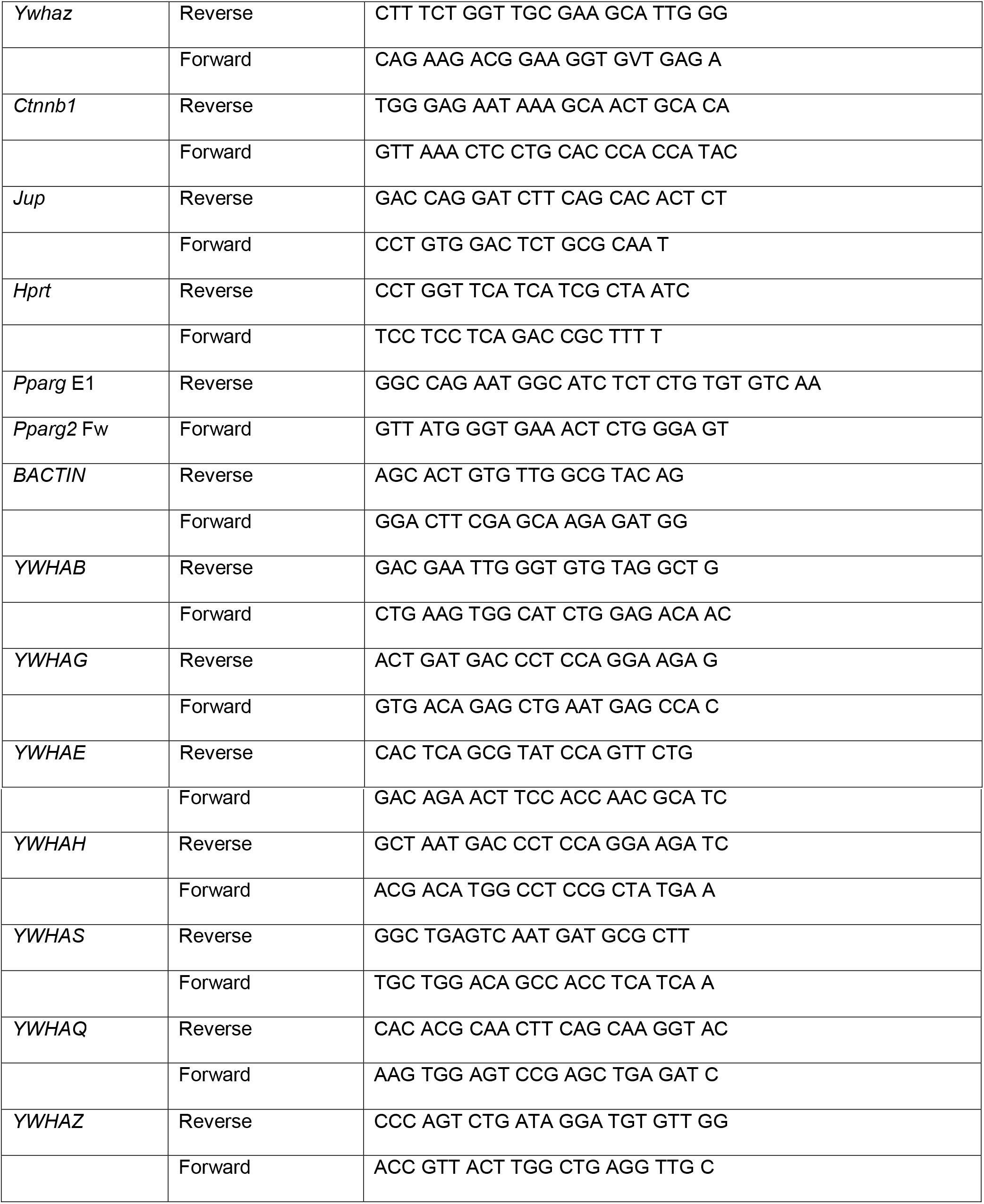
Sequences of primers used in this study.

### Cell fractionation and Western blot

Cell fractionation was used to separate nuclear and cytoplasmic fractions, as previously described (29). In brief 3T3-L1 cells were grown in 100mm dishes, transfected with siRNAs targeting *Ctnnb1* (s63417), *Jup* (s68573), and differentiated for 48 hours before trypsinization and cell collection. Cells were then lysed with Lysis Buffer A (NaCl 150mM, HEPES (pH 7.4) 50nM, Digitonin 25ug/mL, hexylene glycol 1M, protease inhibitor cocktail 1%), and centrifuged at 2000xg for 10 minutes at 4°C to release cytosolic proteins. The supernatant was then collected and Lysis Buffer B (NaCl 150mM, HEPES (pH 7.4) 50nM, Igepal 1% v:v, hexylene glycol 1M, protease inhibitor cocktail 1%) was added, to allow collection of membrane-bound proteins by centrifugation at 7000xg for 10 minutes at 4°C. The supernatant was collected and Lysis Buffer C (NaCl 150mM, HEPES (pH 7.4) 50nM, sodium deoxycholate 0.5% w:v, sodium dodecyl sulfate 0.1% w:v, hexylene glycol 1M, protease inhibitor cocktail 1%) was used to collect nuclear proteins following centrifugation at 7800xg for 10 minutes at 4°C.

For immunoblotting, cells were washed once with PBS and lysed in RIPA buffer (0.9% NaCl, 1% v/v triton X-100, 0.5% sodium deoxycholate, 0.1% SDS, and 0.6% tris base), supplemented with a protease inhibitor (Sigma-Aldrich). Protein lysates were collected, and the concentration was quantified by the Bradford Protein Assay. Proteins were resolved by SDS-PAGE and transferred to PVDF membranes, followed by probing with antibodies for β-catenin (Sigma, #C7207), plakoglobin (Invitrogen, #PG-11E4), 14-3-3ζ (Cell Signaling Technology, #7534) and normalized to β-actin (Cell Signaling Technology, #60). For subcellular fractionation, antibodies for GAPDH (Cell Signaling Technology, #2597) and Lamin A/C (Cell Signaling Technology, #4000) were used. Membranes were visualized with a ThermoFisher Scientific iBright 1500CL, and densitometry was performed with iBrightAnalysis Software (Ver 4.0).

### Data Presentation and Statistical Analysis

Data are presented as mean ± standard error of the mean (SEM) and were visualized with Graphpad Prism (ver 8; San Diego, CA). Statistical significance was calculated by one-way or two-way ANOVA followed by Student’s t-test. The data were considered statistically significant when p < 0.05.

### Data availability

The raw datasets are available as supplementary information in excel sheet formats. Proteomics data has been deposited in ProteomeXchange through partner MassIVE as a complete submission and assigned MSV000088804 and PXD031499. The data can be downloaded from ftp://MSV000088804@massive.ucsd.eduusing the password plako.

## Results

### Determining differences in the 14-3-3ζ interactome in adipose tissue of lean and obese mice

We previously reported that transgenic mice over-expressing a TAP-tagged 14-3-3ζ molecule gained significantly more weight and fat mass when fed a high-fat diet compared to wildtype mice (24). Given the ability of 14-3-3 proteins to interact with a wide array of proteins (21), it raised the possibility that HFD feeding could promote the recruitment of pro-obesogenic factors to the 14-3-3ζ interactome, leading to increased adipogenesis and the expansion of fat mass. As we have previously demonstrated the ability of using proteomics to identify novel regulators of adipocyte differentiation within the 14-3-3ζ interactome (25), we posited that comparing the TAP-14-3-3ζ interactome in gonadal adipose tissue of lean, LFD-fed and obese, HFD-fed mice could lead to the discovery of novel regulators of adipogenesis (Fig. 1A).

**Figure 1.**
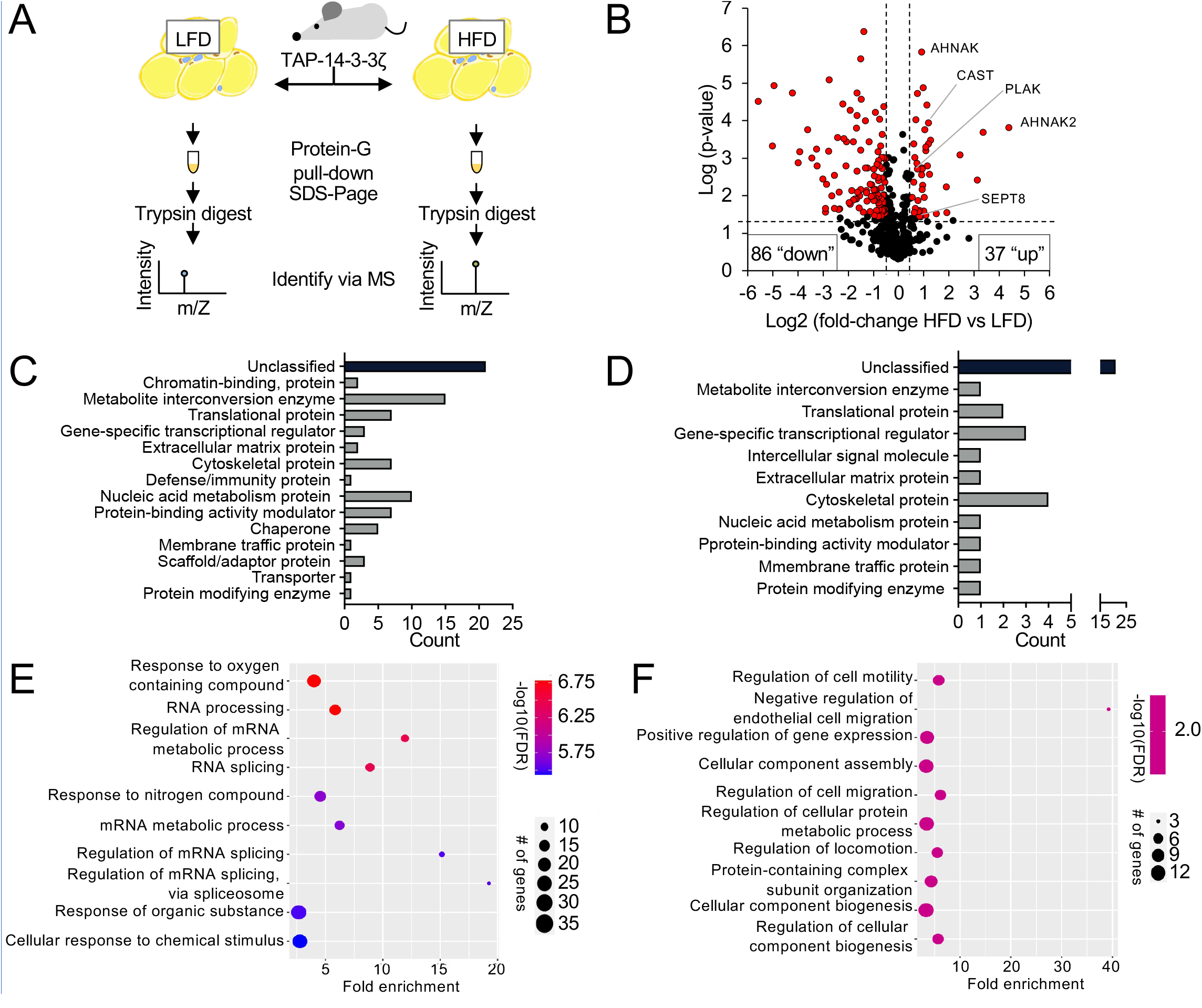
Proteomic analysis reveals differences in the 14-3-3ζ interactome found in gonadal adipose tissue of mice fed a high-fat diet. (**A)** Schematic diagram outlining the workflow to comparing the gonadal adipose 14-3-3ζ interactome between mice fed low-fat (10%, LFD) or high-fat (60%, HFD) diets. (**B)** Visualization of proteins that significantly increased or decreased in abundance in the HFD 14-3-3ζ interactome when compared to the LFD-14-3-3ζ interactome. (**C,D)** Classification by gene ontology of proteins that decreased (C) or increased (D) in abundance according to their putative functions. **(E,F)** Gene ontology was used to categorize proteins that significantly increased (E) or decreased (F) in abundance according to their biological process.

When comparing the TAP-14-3-3ζ interactomes in adipose tissues of LFD- or HFD-fed mice, over 125 unique factors of different proteins classes displayed significantly increased or decreased abundance (Fig. 1B-D; Table S1). Categorization of proteins according to their biological process showed that they were involved in various cellular functions, including protein translation, nucleic acid metabolism, gene transcription, or the cytoskeleton (Fig. 1E,F). Among proteins that were significantly increased in abundance in the HFD-14-3-3ζ interactome, known regulators of adipogenesis, such as Calpastatin (CAST) (30,31) and AHNAK were detected (32,33) (Fig. 1B). Of note was the enrichment of plakoglobin (PLAK), also known as γ-catenin, within the 14-3-3ζ interactome of HFD-fed mice (Fig. 1B). Plakoglobin is closely related to β-catenin, the effector of the Wnt signaling pathways with inhibitory effects on adipocyte differentiation (8–10), but to date, the contributions of plakoglobin to adipocyte differentiation have not been investigated. Knockdown of mRNA corresponding to *Jup*, the gene encoding plakoglobin, in 3T3-L1 pre-adipocytes attenuated the expression of *Pparg* during the first 48 hours of differentiation (Fig. 2A,B). Moreover, depletion of *Jup* mRNA was associated with decreased Oil Red-O incorporation, seven days post-differentiation (Fig. 2C). Consistent with prior studies, depletion of known regulators of adipogenesis, such as CAST and AHNAK, impaired 3T3-L1 differentiation (32,33) (Fig. 2A,C).

**Figure 2.**
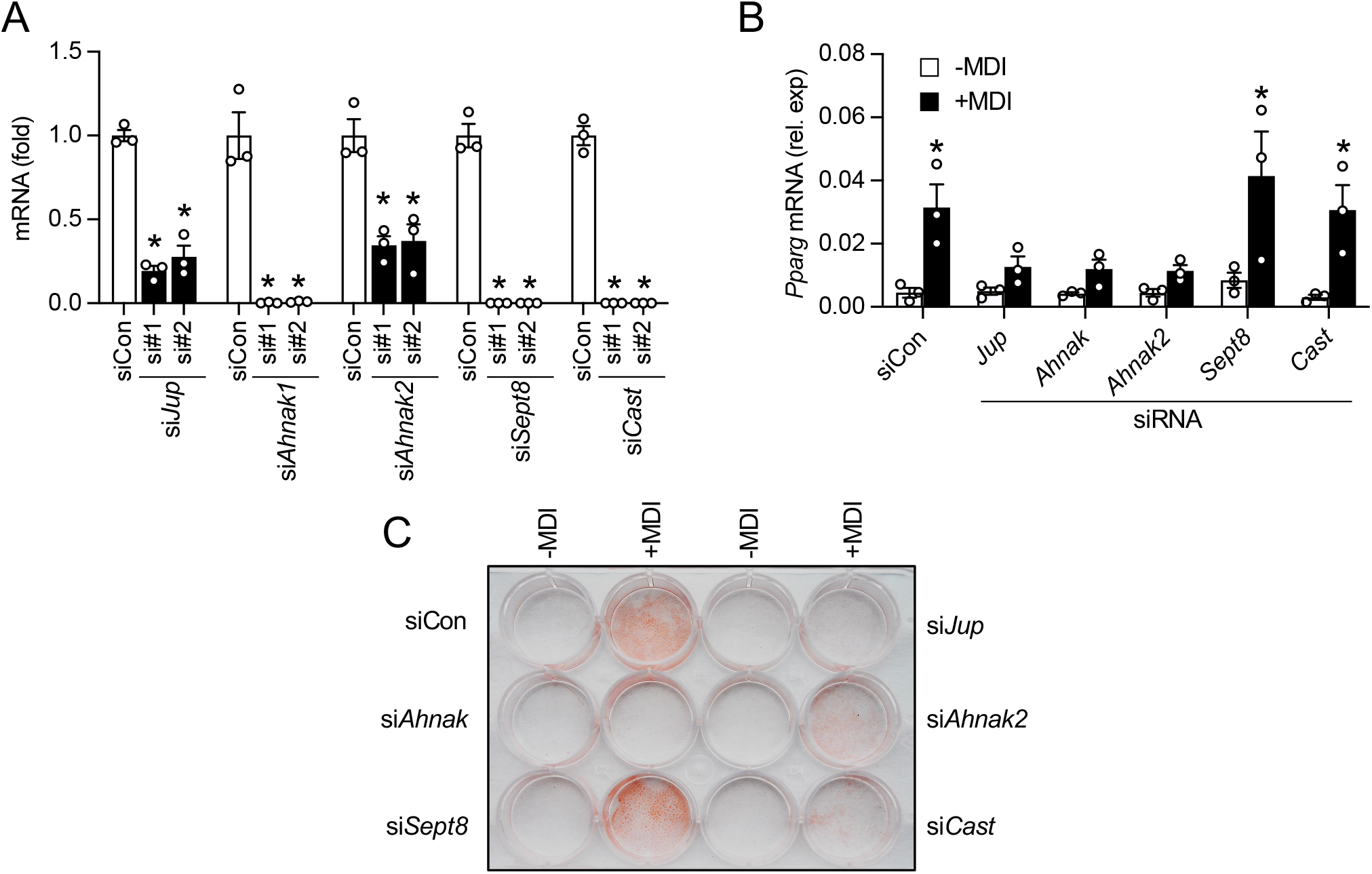
Assessing the role of 14-3-3ζ interactors on adipogenesis. **(A)** Quantitative PCR was used to measure siRNA knockdown efficiency in 3T3-L1 pre-adipocytes. Two independent duplexes were used. All data were normalized to *Hprt*. (n=3 per group, *: p<0.05 when compared to siCon. Significance was determined by one-way ANOVA). **(B)** The role of putative 14-3-3ζ interacting partners in adipogenesis was assessed by siRNA transfection prior to the initiation of adipogenesis (+MDI), followed by RNA isolation after 48 hours and quantitative PCR analysis of *Pparg* mRNA (n=3 per group, *: p<0.05 when compared to the control siRNA, siCon. Significance was determined by two-way ANOVA followed by Bonferroni t-tests). **(C)** Oil Red-O staining was performed on undifferentiated or differentiated 3T3-L1 cells transfected with siRNA against *Ahnak, Ahnak2, Cast, Sept8*, and *Jup* (representative image of n=3 independent experiments). Bar graphs represent mean ± SEM.

### Modulation of plakoglobin or β-catenin expression has opposing effects on PPARγ2 expression during adipogenesis

Given the close homology shared between plakoglobin and β-catenin (34,35), we next sought to compare their relative contributions to adipogenesis. As a first step, 3T3-L1 preadipocytes were transfected with siRNAs targeting 14-3-3ζ, β-catenin and plakoglobin to confirm knockdown. Using two independent siRNAs duplexes, significant reductions in mRNA levels of *Ctnnb1*, the gene encoding β-catenin, and *Jup*, were achieved (Fig. S1A,C). Knockdown of *Ywhaz*, the gene encoding 14-3-3ζ, was used as a positive control (24), and significant reductions in *Ywhaz* mRNA were achieved following siRNA transfection (Fig. S1B). Notably, the induction of adipogenesis significantly decreased *Ctnnb1* mRNA levels in control-transfected (siCon) cells (Fig. S1A). As expected, levels of *Pparg2* mRNA increased following the induction of differentiation (Fig. S1D), and as we previously demonstrated, knockdown of 14-3-3ζ was found to impair *Pparg2* mRNA expression (Fig. S1C) (24,25). Knockdown of *Ctnnb1* and *Jup* downregulated *Pparg2* transcription after 48 hours of the induction of adipogenesis (Fig. S1D).

Immunoblotting was next used to compare protein abundance of PPARγ2 following depletion of β-catenin or plakoglobin. Similar to what was observed with mRNA levels of *Ctnnb1*, β-catenin protein abundance was decreased following the induction of adipogenesis (Fig. 3A), and knockdown of the closely-related homology, plakoglobin, had no effect on β-catenin expression (Fig. 3A). Plakoglobin abundance was significantly decreased following the induction of adipocyte differentiation, and unexpectedly, knockdown of β-catenin led to a 3-fold increase of plakoglobin levels (Fig. 3B). PPARγ2 expression in response to adipogenesis was downregulated by 90 and 69% when 14-3-3ζ or plakoglobin were depleted, respectively (Fig. 3B). The ability of β-catenin depletion has been shown to increase PPARγ2 abundance, resulting in potentiated adipogenesis (36,37), and similarly, β-catenin depletion by siRNA was found to increase PPARγ2 abundance by 1.8-fold (Fig. 3B).

**Figure 3.**
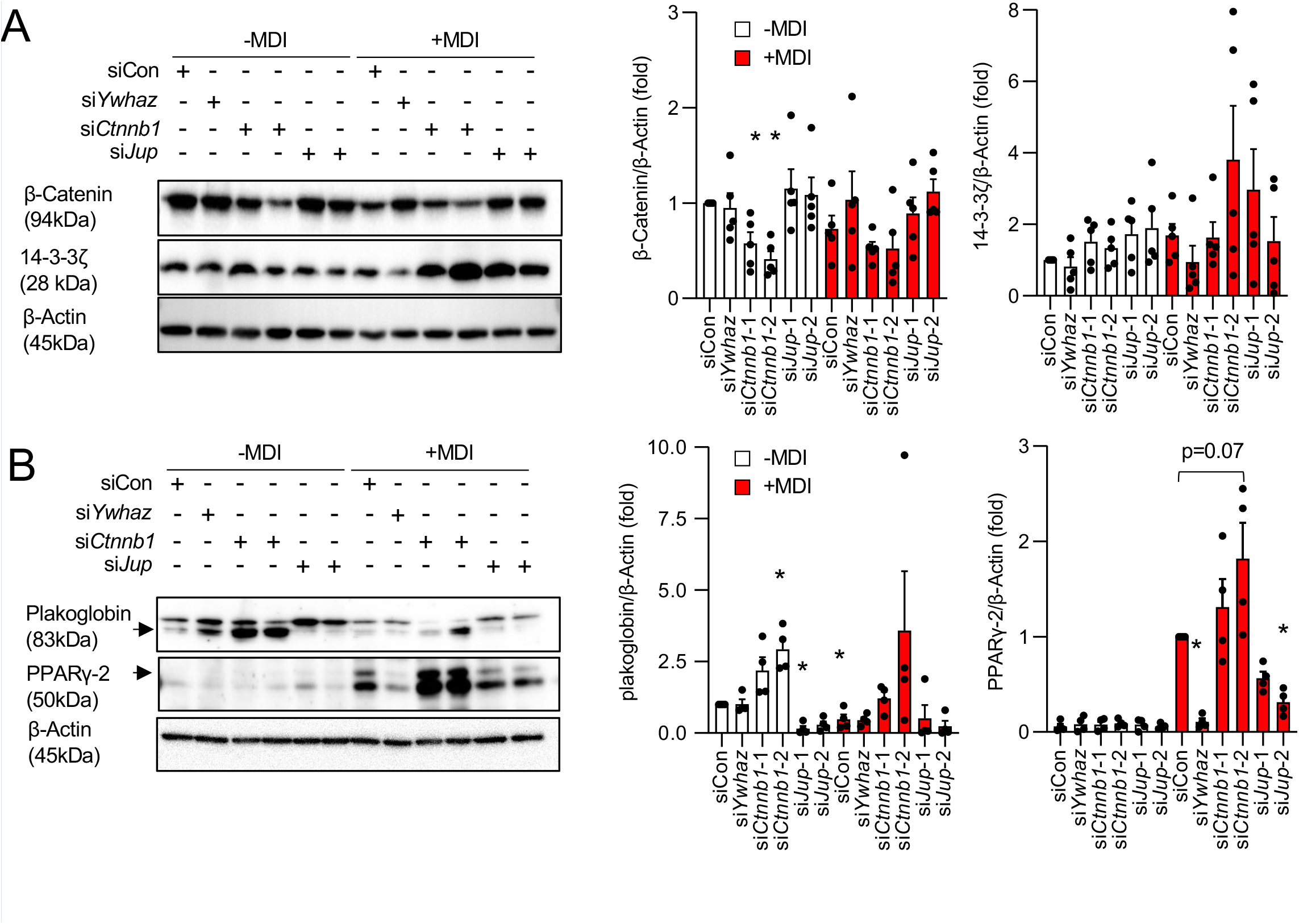
Plakoglobin depletion impairs adipogenesis. **(A)** Protein abundance of β-catenin, β-actin, and 14-3-3ζ in siCon-, *siCtnnb1-, siJup-*, and *siYwhaz*-transfected 3T3-L1 cells prior to and 48 hours after the induction of differentiation (+MDI). (**B)** Protein abundance of plakoglobin, β-actin, and PPARγ2 in siCon-, *siCtnnb1-, siJup-*, and *siYwhaz*-transfected 3T3-L1 cells prior to and 48 hours after the induction of differentiation. Bar graphs represent mean ± SEM. (n=4-5 per group; *: p<0.05 when compared to siCon,-MDI; Significance was determined by two-way ANOVA and student t-test).

We next evaluated the effects of transient overexpression of plakoglobin and β-catenin in 3T3-L1 preadipocytes. Previous studies have reported β-catenin’s ability to inhibit adipogenesis when stably expressed in 3T3-L1 cells, as well as in adipocyte progenitors (10,38). This resulted in adipose tissue fibrosis (38) and commitment of mesenchymal stem cells towards osteogenesis (39,40). In agreement with previous findings, overexpression of β-catenin downregulated PPARγ2 abundance by 34% (p ≤.0.05) following 48 hours of MDI treatment (Fig. 4A). Unexpectedly, plakoglobin overexpression further diminished PPARγ2 expression by 63% (p ≤.0.05) compared to control 3T3-L1 cells (Fig. 4B). When taken together, these findings demonstrate that plakoglobin and β-catenin can have similar and divergent roles in adipogenesis depending on their relative expression levels.

**Figure 4.**
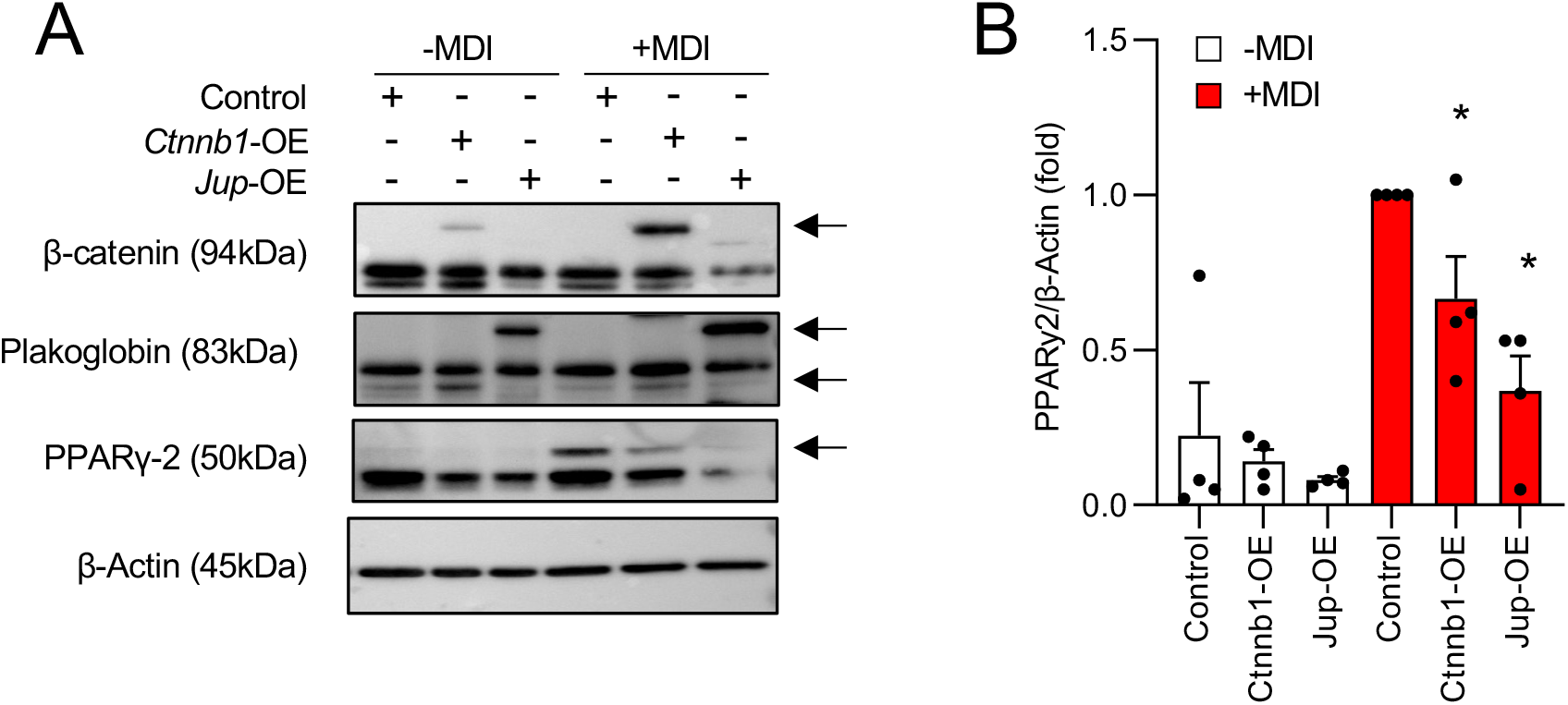
Plakoglobin overexpression impairs adipogenesis. **(A)** Protein abundance of β-catenin, plakoglobin, β-actin, and PPARγ2 in 3T3-L1 transiently transfected were treated with plasmids encoding mEmerald-tagged plakoglobin or β-catenin, followed by induction of adipogenesis (+MDI) (representative of n=4 independent experiments). **(B)** Quantification of PPARγ2 abundance following ectopic overexpression of plakoglobin or β-catenin in 3T3-L1 cells. All data were normalized to β-actin. (n=4 per group; *: p<0.05 when compared to siCon+MDI; Significance was determined by two-way ANOVA and student t-test). Bar graphs represent mean ± SEM.

### Nuclear localization of plakoglobin and β-Catenin are decreased during adipogenesis

Plakoglobin and β-catenin bind to TCF transcription factors, such as TCF7L2 (41,42), which facilitates their nuclear translocation. As depletion of plakoglobin and β-catenin have opposing effects on the expression of PPARγ2 during adipogenesis (Fig. 5A), we next examined whether β-catenin or plakoglobin exhibit differences in nuclear abundance during adipogenesis to influence gene transcription. Nuclear fractionation was first used to assess the localization of both proteins in naïve and differentiating 3T3-L1 cells. β-catenin was primarily localized to the nucleus, and its abundance in the nuclear compartment was unaffected by plakoglobin depletion. Following differentiation, nuclear β-catenin abundance decreased (Fig. 5A). Similar to β-catenin, plakoglobin was primarily restricted to the nucleus and protein abundance markedly decreased following the induction of differentiation (Fig 5B). Plakoglobin nuclear abundance was markedly upregulated following siRNA-mediated depletion of β-catenin, but this increase in abundance was not maintained in 3T3-L1 cells undergoing differentiation (Fig. 5B).

**Figure 5.**
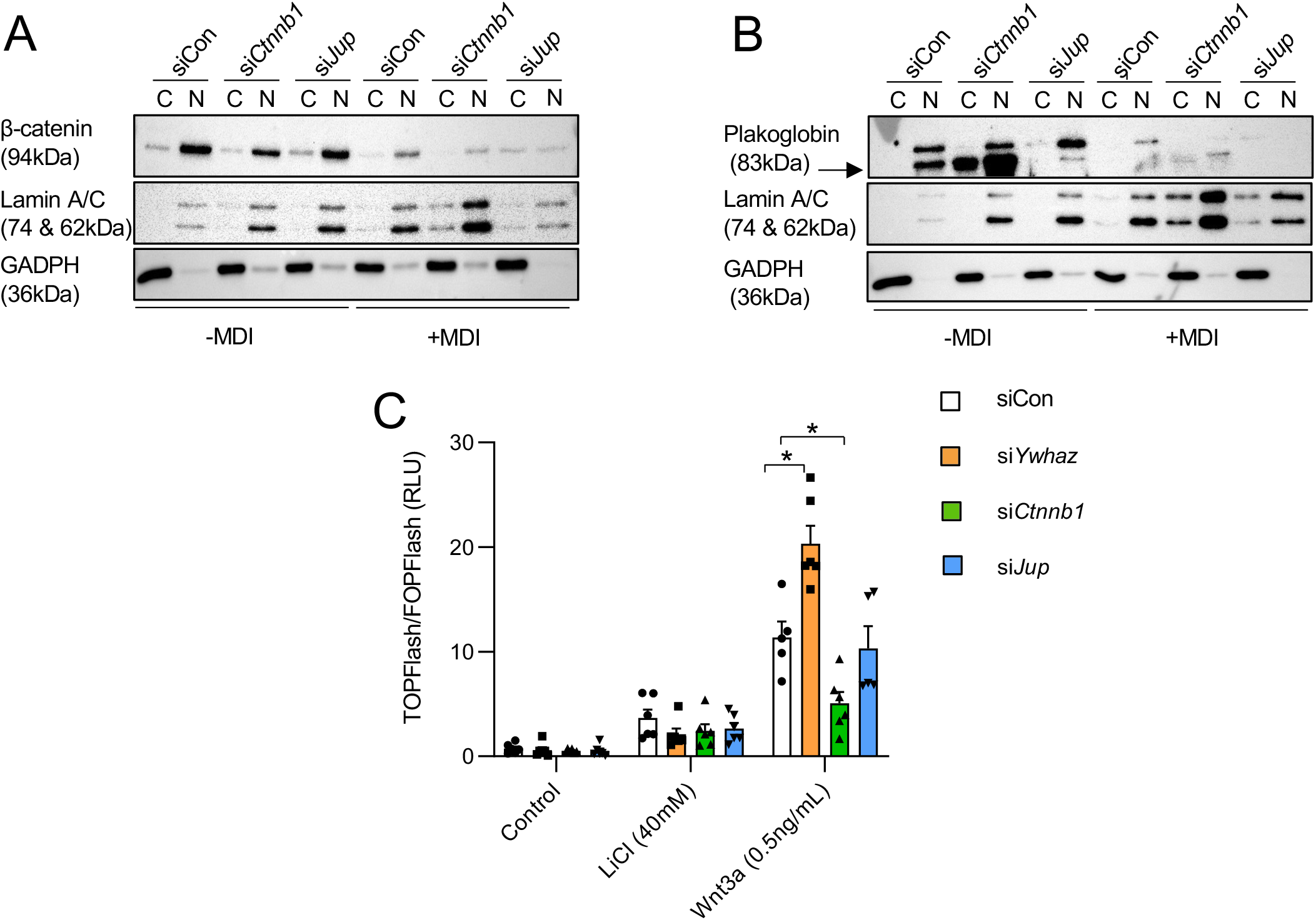
Plakoglobin is exclusively found in the nucleus of 3T3-L1 cells and does not participate in Wnt-induce transcriptional activity. **(A,B**) Immunoblotting was used to measure protein abundance of β-catenin (A) and plakoglobin (B) in the cytoplasm and nucleus of 3T3-L1 cells that were induced to differentiate (+MDI) for 48 hours. Cells were transfected with either siCon, si*Jup*, or *siCtnnb1*. Lamin A/C or GAPDH were used as nuclear or cytoplasmic markers, respectively (representative of n=4 independent experiments). **C.** Wnt/β-catenin transcriptional activity as assessed by SuperTOPFlash following transfected with *siYwhaz, siCtnnb1*, and *siJup*. All data were normalized to cells transfected with FOPFlash. Activation of the Wnt pathway was induced by LiCl (40mM) and Wnt3a (0.5ng/mL). (n=5-6 per group; *\**: p<0.05 when compared to siCon; Significance was determined by one-way ANOVA within treatment group). Bar graphs represent mean ± SEM.

As depletion of β-catenin increased plakoglobin nuclear abundance, we next examined whether this would lead to changes in Wnt transcriptional activity. NIH-3T3 cells were transfected with SuperTOPFlash and FOPFlash reporter plasmids, followed by transfection with siRNA against 14-3-3ζ, β-catenin, or plakoglobin. Transfected cells were then treated with 40mM LiCl or 0.5ng/mL Wnt3a for 24 hours. In LiCl-treated cells, a 4.11-fold increase in transcriptional activity was observed in siCon-transfected cells, and knockdown of 14-3-3ζ, β-catenin, or plakoglobin had no effect. Treatment with Wnt3a was associated with a 12-fold increase (p<0.05) in transcriptional activity in siCon-transfected cells (Fig. 5C). 14-3-3ζ is known to interact with β-catenin to limit its nuclear localization, suggesting a regulatory role on Wnt transcriptional activity (43). Indeed, depletion of 14-3-3ζ potentiated Wnt3a-induced increased transcriptional activity and demonstrated the ability of 14-3-3ζ to function as a negative regulator of the Wnt signaling pathway. Despite the high degree of homology, depletion of plakoglobin had no effect on Wnt3a-induced transcriptional activity (Fig. 5C), which suggests that plakoglobin is not involved in the regulation of Wnt3a-associated transcriptional activity.

### Depletion of β-catenin does not fully rescue the effect of plakoglobin knockdown on adipogenesis

As β-catenin depletion increased PPARγ2 abundance under differentiating conditions (Fig. 3B), we next sought to examine whether β-catenin depletion could rescue plakoglobin-deficient 3T3-L1 pre-adipocytes from defective adipogenesis. Quantitative PCR was first used to confirm effective concomitant knockdown of *Ctnnb1* and *Jup* mRNA (Fig. S2A,B). Levels of *Pparg2* mRNA were significantly increased in *Ctnnb1*-depleted, undifferentiated cells and also in double knockdown, undifferentiated cells (Fig. 6A-inset), and following the induction of differentiation, *Pparg2* mRNA levels were significantly higher in *siCtnnb1*- and *siCtnnb1* + si*Jup*-transfected cells when compared to siCon-transfected cells (Fig. 6A).

**Figure 6.**
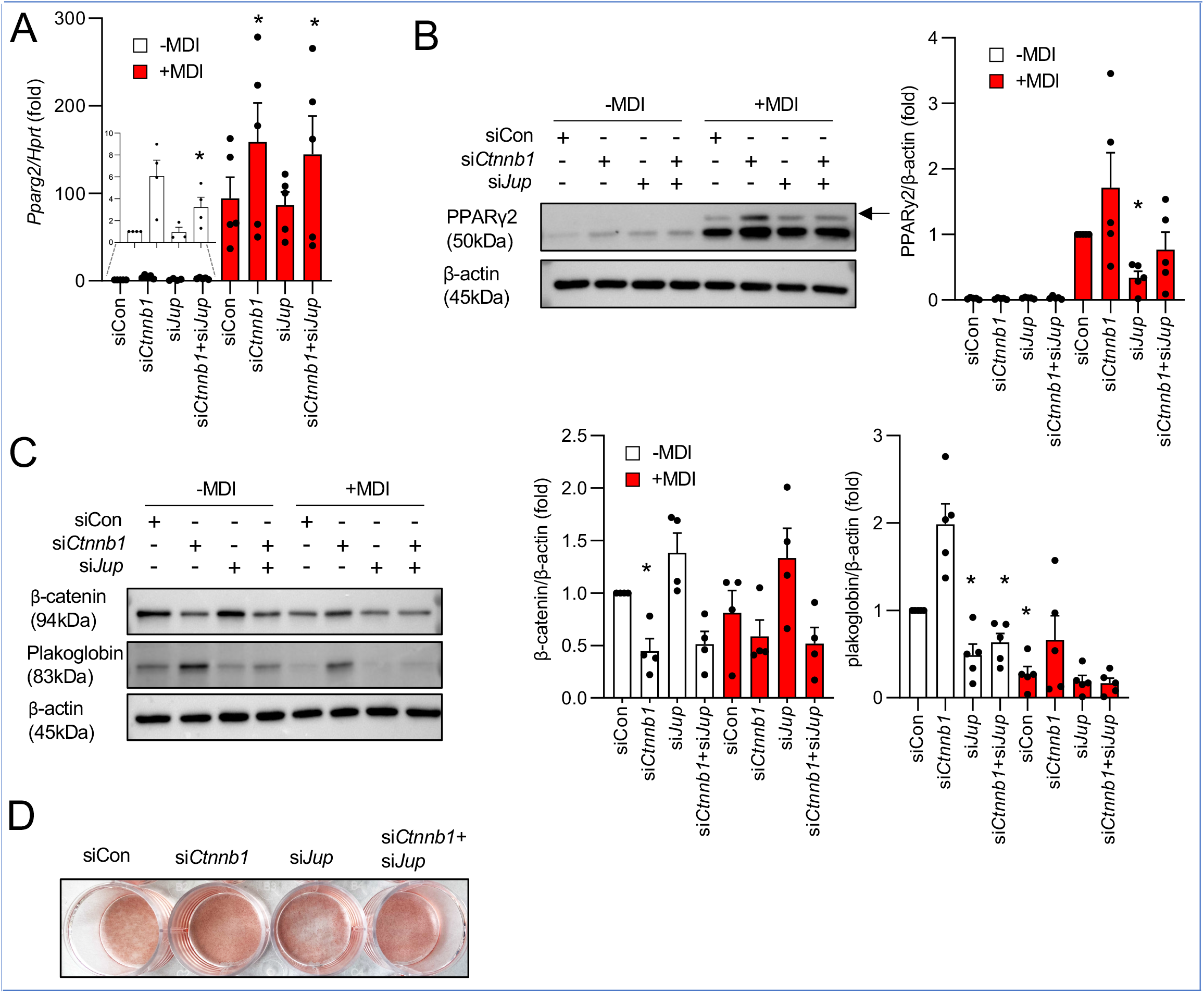
Double knockdown of plakoglobin and β-Catenin increases *Pparg2* expression but does not affect PPARγ2 abundance. **(A)** Levels of *Pparg2* mRNA following single or double knockdown of *Jup* or *Ctnnb1* and in response the induction of adipogenesis (+MDI). The inset image depicts *Pparg* mRNA levels in undifferentiated cells (n=5 per group; *: p<0.05 when compared to siCon-MDI; Significance was determined by two-way ANOVA and student t-test). **(B)** Immunoblotting was used to measure protein abundance of PPARγ2 in undifferentiated and differentiating 3T3-L1 cells transfected with siRNA against *Ctnnb1* or *Jup* (n=4-5 per group; *: p<0.05 when compared to siCon+MDI; Significance was determined by two-way ANOVA and student t-test). **(C)** Measurement of β-catenin and plakoglobin plakoglobin in single- or double-transfected 3T3-L1 cells induced to differentiate (+MDI) (n=4-5 per group; *: p<0.05 when compared to siCon-MDI; Significance was determined by two-way ANOVA and student t-test). **(D)** Oil Red-O staining of fully differentiated 3T3-L1 cells transfected with siRNA targeting β-catenin, plakoglobin, or both (representative images of n=4 independent experiments). Bar graphs represent mean ± SEM.

Although single depletion of β-catenin was found to upregulate plakoglobin expression (Fig. 4B), concomitant knockdown was unable to prevent the loss of plakoglobin due to siRNA-mediated depletion, suggesting that β-catenin’s influence on plakoglobin expression is transcriptionally regulated (Fig. 6C,S2B). Immunoblotting for PPARγ2 revealed that the concomitant depletion of β-catenin or plakoglobin was sufficient to increase PPARγ2 protein abundance in differentiating cells (Fig. 6B). Measurement of lipid content by ORO incorporation after a seven day differentiation period revealed that single knockdown of β-catenin and plakoglobin were associated with enhanced and impaired ORO incorporation, respectively (Fig. 6D). Double knockdown of both β-catenin and plakoglobin was associated with ORO incorporation similar to siCon-transfected cells (Fig. 6D).

### Effects of plakoglobin or β-catenin depletion on human adipogenesis

Of the seven mammalian 14-3-3 isoforms we have shown that 14-3-3ζ is required for murine adipogenesis *in vitro* and *in vivo* (24), but it is not clear whether 14-3-3ζ is required for human adipocyte differentiation. To this end, siRNAs against *YWHAB, YWHAE, YWHAG, YWHAH, YWHAQ, YWHAS*, and *YWHAZ* were transfected into hADSCs prior to differentiation followed by analysis of lipid content by Oil Red-O after 14 days of differentiation. Quantitative PCR confirmed effective knockdown of each isoform (Fig, 7A). Of the seven human isoforms, Oil Red-O incorporation revealed that 14-3-3σ and 14-3-3ζ were required for human adipogenesis (Fig. 7B). We next assessed the effects of β-catenin and plakoglobin depletion on hADSC differentiation. Knockdown of either β-catenin or plakoglobin was successfully attained and confirmed by immunoblotting (Fig. 7C,D). Comparable to 3T3-L1 cells, β-catenin knockdown enhanced lipid accumulation (Fig. 6D,7E). Similarly, depletion of plakoglobin led to reduce lipid content, suggesting defects in adipogenesis (Fig, 7E).

**Figure 7.**
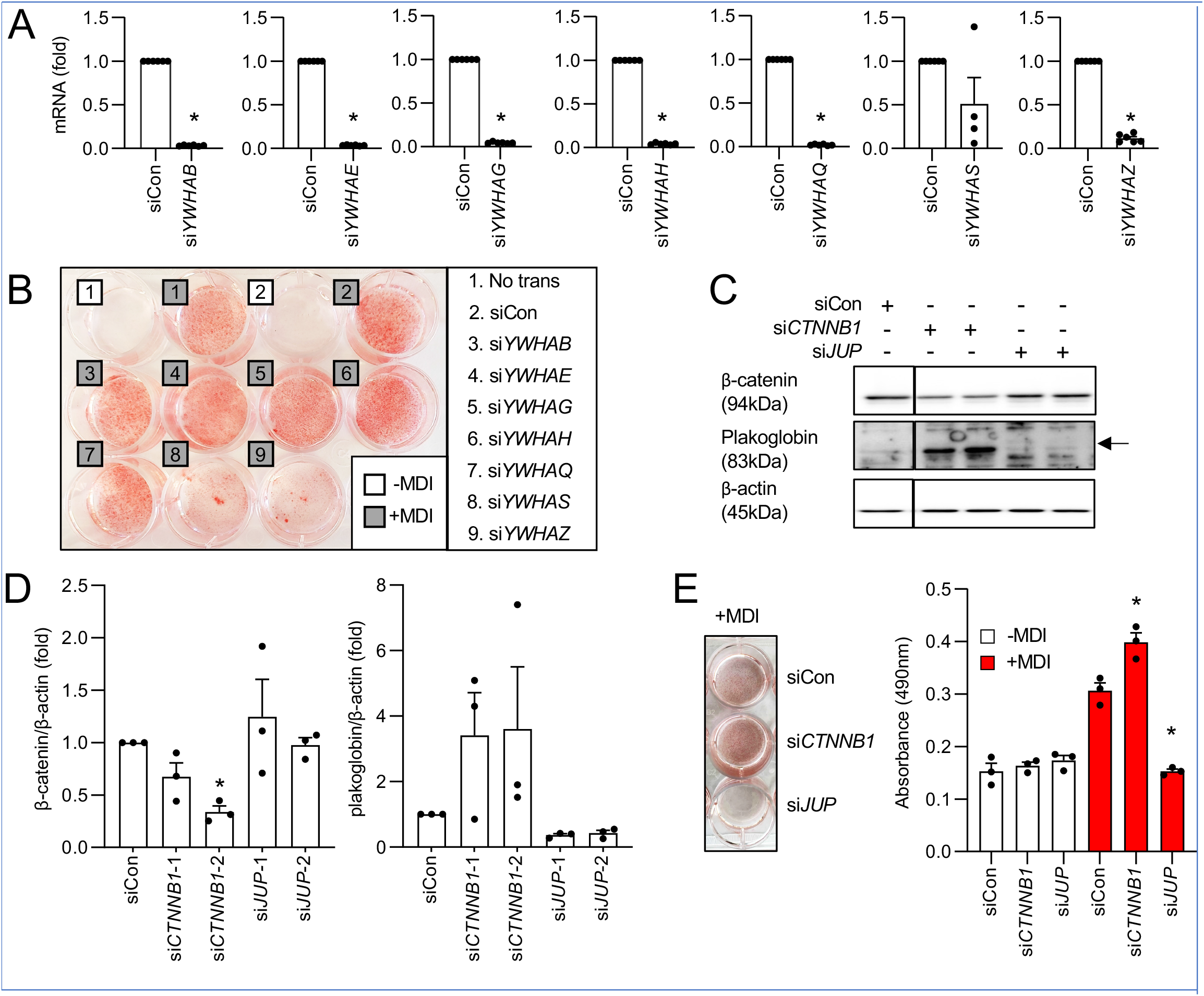
14-3-3ζ, 14-3-3σ, and plakoglobin are required for the differentiation of human adipocyte stem cells (hADSCs). **(A)** Quantitative PCR was used to assess knockdown efficiency of siRNA duplexes against human 14-3-3 mRNA isoforms. Data were normalized to *ACTB* mRNA (n=4 per group; *: p<0.05; Significance was determined by Student’s t-test). **(B)** Oil Red-O staining was used to determine adipocyte differentiation (+MDI) following transfection with siRNA targeting all human 14-3-3 isoforms (representative of n=4 independent experiments). **(C,D)** Immunoblotting (C) and densitometric quantification (D) of β-catenin and plakoglobin abundance in hADSCs following siRNA-mediated depletion of *CTNNB1* or *JUP*. **(E)** Oil Red-O staining was used to assess hADSCs differentiation in cells transfected with *siJUP* or *siCTNNB1* prior to differentiaton (+MDI). (n=3 per group; *: p<0.05 when compared to siCon+MDI; Significance was determined by one-way ANOVA). Bar graphs represent mean ± SEM.

## Discussion

The signaling events underlying adipogenesis have been extensively studied, but a complete understanding of the many adipogenic regulators is lacking. Our study shows that the interactome of 14-3-3ζ can be successfully mined to discover novel, adipogenic regulators, such as β-catenin’s close homolog, plakoglobin. While β-catenin is known to inhibit adipogenesis, plakoglobin’s influence on adipocyte differentiation has yet to be studied in detail. Herein, we demonstrate a novel role of plakoglobin in adipogenesis. In 3T3-L1 pre-adipocytes, plakoglobin is uniquely found in the nucleus, where it may influence the transcription of downstream target genes, and following the induction of adipogenesis, its abundance is downregulated. Depletion of plakoglobin did not affect Wnt3a-stimulated β-catenin signalling activity, implying that plakoglobin does not participate in Wnt-dependent transcriptional activity. With the successful identification and validation of plakoglobin as a regulator of adipogenesis, our study demonstrates the usefulness of elucidating the 14-3-3ζ interactome to under disease conditions, such as obesity, to identify factors that may contribute to disease development.

Previous work has highlighted the usefulness of identifying how the 14-3-3ζ interactome changes in response to stimuli, as it permits an understanding of how 14-3-3ζ may influence cellular processes (44–46). For example, we previously showed that the 14-3-3ζ interactome becomes enriched with various RNA splicing factors during adipocyte differentiation and that the presence of 14-3-3ζ was necessary for RNA splicing (25). Using a similar proteomic approach, we now show that the interactome of 14-3-3ζ within murine gonadal adipose tissue can be influenced by high-fat diet feeding and the development of obesity. Of the 125 unique factors that significantly changed in abundance, known factors required for adipogenesis or high-fat diet induced obesity, such as AHNAK and Calpastatin, were detected (30–33). It should be noted that it is not clear whether 14-3-3ζ directly interacts with either AHNAK or Calpastatin or whether it influences theirs activities, but this represents an important future aspect to explore.

The identification of plakoglobin in the HFD-14-3-3ζ interactome was unexpected, and given the close homology and reported over-lapping actions with β-catenin (18,34,35), it prompted further examination to determine its contributions to adipogenesis. Knockout models in cardiomyocytes have indicated that plakoglobin is responsible for the increased presence of adipocytes seen in cardiac tissue of mice afflicted by arrhythmic right ventricular cardiomyopathy (19). Recently, plakoglobin’s involvement in insulin signalling and glucose uptake was discovered in mature 3T3-L1 adipocytes (49), but no studies have been performed within preadipocytes. In the present study, depletion of plakoglobin in 3T3-L1 pre-adipocytes was sufficient to reduce *Pparg2* expression and significantly diminish PPARγ2 abundance. PPARγ2 is indispensable for adipogenesis and is responsible for initiating the transcription of multiple genes involved in lipid metabolism (2,50). Surprisingly, plakoglobin overexpression was found to downregulate PPARγ2 expression levels in response to the induction of adipogenesis. The mechanism accounting the inhibitory effect of plakoglobin over-expression is not clear, but since PPARγ2 has a catenin-binding site, high levels of nuclear plakoglobin could result in interactions with PPARγ2 and alterations in its pro-adipogenic activity (51).

Adipogenesis is a two-step process, where the first phase requires the commitment of mesenchymal stem cells into pre-adipocytes, followed by pre-adipocytes terminally differentiating into adipocytes (2). As 3T3-L1 cells are committed pre-adipocytes, the inhibition of adipogenesis following β-catenin or plakoglobin deletion suggests that they are critical for controlling the expression of the first wave of transcription factors, that lie downstream of the commitment phase of mesenchymal stem cells (52). Indeed, our investigation in human adipocyte stem cells supports this hypothesis.

We found that the induction of adipogenesis leads to the degradation of both plakoglobin and β-catenin within the nucleus. While it has been reported that PPARγ2 may facilitate β-catenin degradation during adipogenesis (51), it is not clear whether plakoglobin is similarly regulated. Nonetheless, it is possible that both plakoglobin and β-catenin stability may be controlled by the same degradation complex (53,54). Plakoglobin and β-catenin are known to competitively bind to TCF7L2 in a context-dependent manner, thereby affecting the ability of TCF7L2 to bind DNA and initiate the gene transcription (41). Our data suggests only β-catenin, but not plakoglobin, affects TCF-mediated transcriptional activity, as only β-catenin depletion abrogated Wnt3a-stimulated SUPERTOPFlash activity. The molecular mechanisms that govern the nuclear translocation of β-catenin and plakoglobin have not been studied extensively, but 14-3-3ζ has been shown to restrict its localization to the cytoplasm (43). Indeed, our finding that 14-3-3ζ depletion potentiated Wnt3A-stimulated transcriptional activity supports the idea that 14-3-3ζ may have inhibitory effects on Wnt-β-catenin signaling. Thus, while plakoglobin and β-catenin may share common regulatory pathways or mechanisms, their functions in Wnt-mediated activity can differ.

Human adipocyte stem cells are an established culture model for studying adipogenesis (55–57), and we now show that they are a viable cellular model to examine the roles of 14-3-3 proteins in adipogenesis. Similar to our previous study (24), we now report that 14-3-3ζ is also necessary for human adipogenesis. It should be noted that 14-3-3σ was also found to be required for human adipocyte stem cell differentiation, which is in contrast to what we reported in murine 3T3-L1 cells pre-adipocytes. Consistent with our data with 3T3-L1 cells, plakoglobin is also required for human adipocyte differentiation, as its depletion lead to impaired lipid acculmulation. This further highlights the importance of plakoglobin in the regulation of adipogenesis.

In conclusion, the current study demonstrates that plakoglobin holds a key role in regulating adipocyte differentiation. Moreover, it places particular emphasis on the usefulness of using proteomics to uncover novel adipogenic regulators within the 14-3-3ζ interactome. Plakoglobin is required for the adipogenic differentiation *in vitro*, and the mechanism by which plakoglobin influences adipogenesis appears to be independent of the Wnt/β-catenin signaling pathways. While additional studies aimed at assessing the *in vivo* contributions of plakoglobin to adipocyte differentiation and the expansion of adipose tissue mass are required, our novel findings further expand and deepen our knowledge of adipogenic regulators that could contribute to the development of obesity.

## Financial support

This work is supported by CIHR Project (PJT-153144) and a Discovery Award from the Banting Research Foundation. This work was also funded by CIHR Project (PJT-152995 to P.P.R). GEL holds the Canada Research Chair in Adipocyte Development. SDV was supported by a IUBMB Wood-Whelan Fellowship. PPR is a Senior Scholar of the Fonds de la recherche du Québec-Santé (FRQS). Proteomic analyses were performed by the Center for Advanced Proteomics Analyses (CAPCA), a Node of the Canadian Genomic Innovation Network that is supported by the Canadian Government through Genome Canada.

## Contributions

FA and GEL wrote the manuscript. FA, YM, SY, SDV, GL, PPR, GEL performed experiments, analyzed data, and edited the manuscript. GEL edited the manuscript and the guarantor of this work.

## Conflict of interest

The authors declare that they have no conflicts of interest with the contents of this article.

## Notes

### Competing Interest Statement

The authors have declared no competing interest.

