## Supplemental figures for "Plakoglobin regulates adipocyte differentiation independently of the Wnt/β-catenin signalling pathway"

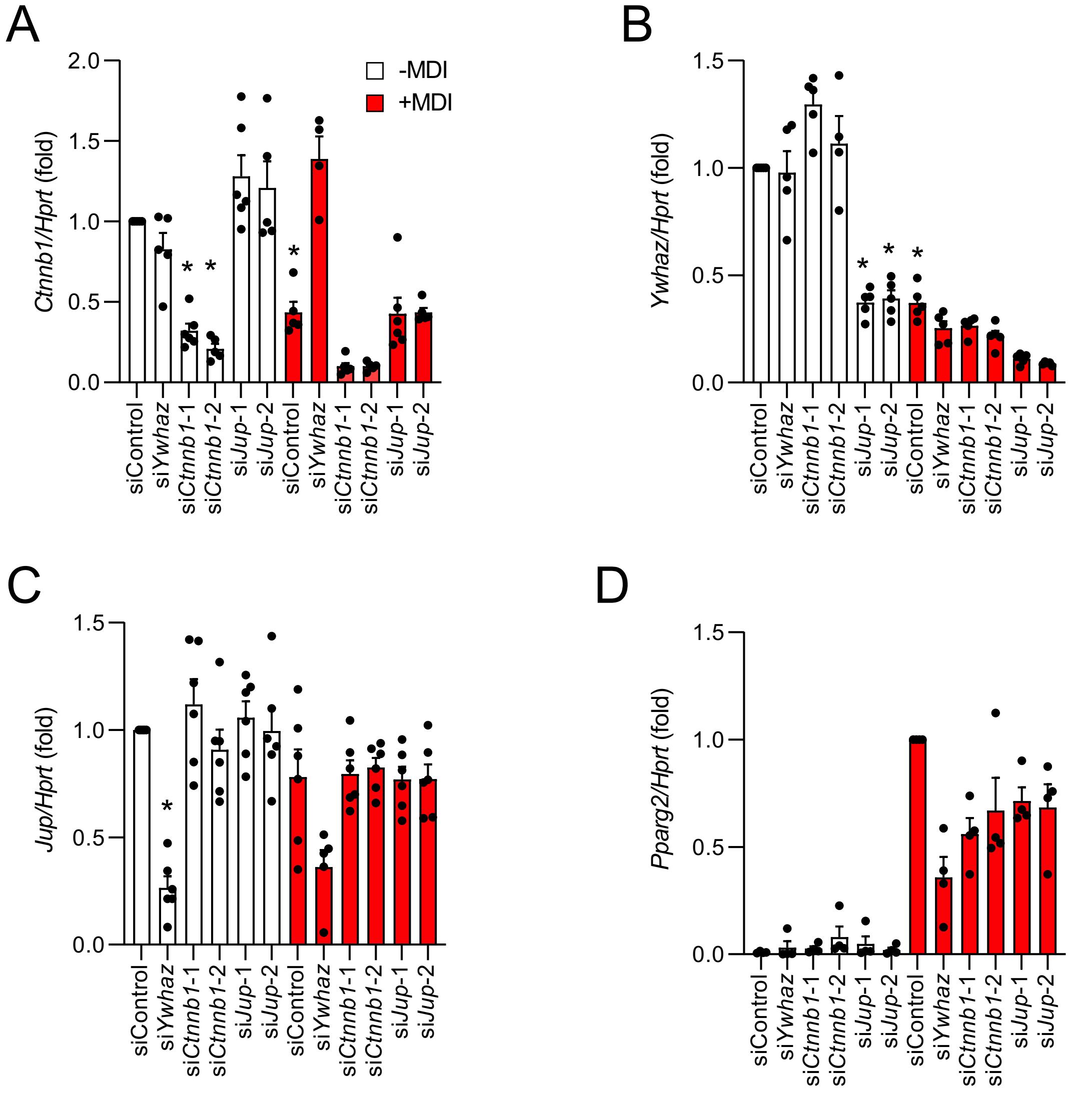


**Figure S1. *Jup, Ctnnb1, and Ywhaz* depletion downregulate *Pparg2* expression in differentiating adipocytes.** **A-D.** Gene expression of *Ctnnb1* **(A)**, *Jup* **(B)**, *Ywhaz* **(C)** and *Pparg2* **(D)** following siRNA mediated depletion and adipogenic induction (n=4-6 per group; *: $p<0.05$; Significance determined by two-way ANOVA and student t-test). Bar graphs represent mean ± SEM.

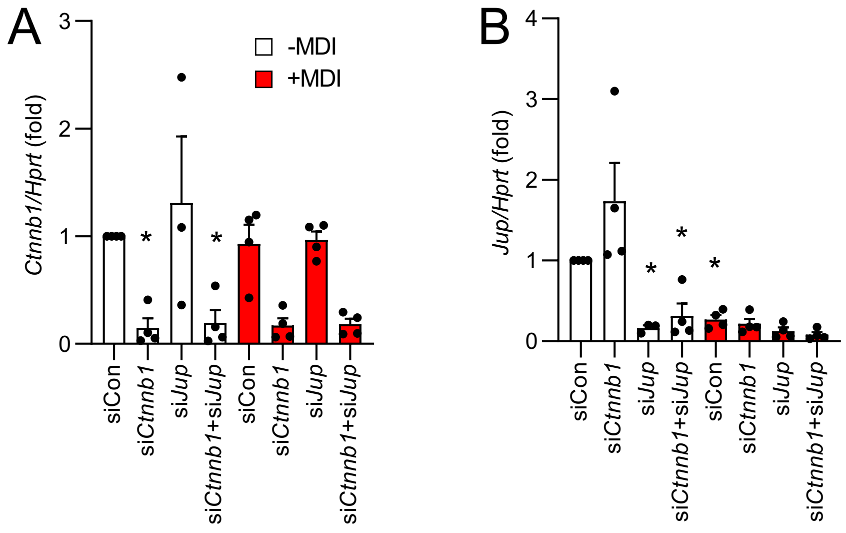


**Figure S2. Quantitative PCR was used to assess *Jup* and *Ctnnb1* mRNA levels in 3T3-L1 cells singly or doubly transfected with siRNA against *Jup* or *Ctnnb1.*** Levels of *Ctnnb1* (**A**) or *Jup* (**B**) mRNA, as assessed by quantitative PCR, in 3T3-L1 adipocytes singly or doubly transfected with si*Ctnnb1* or si*Jup*. Expression of either gene was examined in undifferentiated (-MDI) or cells induced to differentiate for 48 hours (+MDI) (n=4 per group; *: $p<0.05$ when compared to siCon-MDI; Significance was determined by two-way ANOVA and Bonferroni t-tests). Bar graphs represent mean ± SEM.
